# Protein structural disorder of the envelope v3 loop contributes to the switch in human immunodeficiency virus type 1 cell tropism

**DOI:** 10.1101/130161

**Authors:** Xiaowei Jiang, Felix Feyertag, David L. Robertson

## Abstract

Human immunodeficiency virus type 1 (HIV-1) envelope gp120 is partly an intrinsically disordered (unstructured/disordered) protein as it contains regions that do not fold into well-defined protein structures. These disordered regions play important roles in HIV’s life cycle, particularly, V3 loop-dependent cell entry, which determines how the virus uses two coreceptors on immune cells, the chemokine receptors CCR5 (R5), CXCR4 (X4) or both (R5X4 virus). Most infecting HIV-1 variants utilise CCR5, while a switch to CXCR4-use occurs in the majority of infections. Why does this ‘rewiring’ event occur in HIV-1 infected patients? As changes in the charge of the V3 loop are associated with this receptor switch and it has been suggested that charged residues promote structure disorder, we hypothesise that the intrinsic disorder of the V3 loop plays a role in determining cell tropism. To test this we use three independent data sets of gp120 to analyse V3 loop disorder. We find that the V3 loop of X4 virus has significantly higher intrinsic disorder tendency than R5 and R5X4 virus, while R5X4 virus has the lowest. These results indicate that structural disorder plays an important role in determining HIV-1 cell tropism and CXCR4 binding. We speculate that changes in N-linked glycosylation associated with tropism change (from R5 to X4) are required to stabilise the V3 loop with increased disorder tendency during HIV-1 evolution. We discuss the potential evolutionary mechanisms leading to the fixation of disorder promoting mutations and the adaptive potential of protein structural disorder in viral host adaptation.

**IMPORTANCE:** HIV-1 cell entry relies on the V3 loop of its heavily glycosylated envelope protein gp120 to bind to a host coreceptor CCR5 or CXCR4. Unraveling the mechanism whereby HIV-1 switches host coreceptor is critical to understanding HIV-1 pathogenesis and development of novel intervention strategies. However, a mechanistic understanding of the switch is limited as no gp120-CCR5/CXCR4 complex is available, due to the intrinsically disordered nature of the V3 loop responsible for coreceotor swtich. We hypothesise that shifts of V3 disorder may contribute to HIV-1 coreceptor switch and cell tropism. In this study we compared the disorder tendency of the V3 loop before and after the coreceptor switch. We find that the coreceptor switch is associated with a significant increase of V3 loop disorder from CCR5 to CXCR4 using. This result provides a mechanistic explanation of coreceptor switch that increasingly disordered V3 loop results in use of a different host coreceptor.

## Introduction

HIV-1 cell entry relies primarily on the binding of viral envelope protein, gp120, to two host immune cell membrane proteins, namely, the CD4 receptor and the coreceptor CCR5 or CXCR4 (1). Virus that solely uses CCR5 as coreceptor is termed R5 tropic, whereas exclusive CXCR4 using virus is termed X4 tropic, while virus that can use both coreceptors for cell entry is termed dual tropic (R5X4). The switch from CCR5 to CXCR4 is hypothesised to play an important role in disease progression and pathogenesis (2, 3). Interestingly, the CCR5 receptor is not expressed in individuals that present the CCR5 delta-32 mutation (4). This mutation confers natural resistance to R5 tropic HIV and non-expression of this receptor appears not to have any significant affect. Based on these observations, the small molecule CCR5 antagonist maraviroc (MVC) was developed to inhibit the CCR5 receptor, thereby blocking virus cell entry (5) and helping to control virus infection.

In all documented cases of MVC virologic failure due to an observed tropism switch, a minority pre-existing X4 using viral population was present prior to therapy, which is ‘unmasked’ by the use of an entry-inhibitor (6). This indicates that X4 populations may commonly be present, but only transiently dominant, in the life time of an infection. Indeed, in the now classic longitudinal study of Shankarrapa et al. (7), this is readily apparent (see Meehan et al. for visualisations (8)). This indicates that the true rates of X4 using virus are much higher than the commonly reported figure that 50% of patients are observed to progress to X4 using virus, as confirmed by recent studies (3, 9).

Investigating HIV-1 coreceptor switch is therefore central to understanding HIV-1 infection and has clinical implications for the use of CCR5 entry inhibitors. It is generally accepted that the V3 loop of gp120 plays a primary role in determining HIV-1 cell tropism and coreceptor specificity (10). However, our understanding of the mechanisms of HIV-1 tropism remains incomplete (2, 3). Charged residues in the V3 loop have been shown to affect tropism (11, 12). Moreover, N-linked glycosylation in the vicinity of the V3 loop has been shown to affect viral tropism(13), e.g., the lack of N-linked glycosylation sites is associated with X4 phenotypes (13-18). Because N-linked glycosylation plays a role in protein folding and stability (19-22), lack of N-linked glycosylation may decrease V3 loop stability and therefore contribute to X4 using phenotypes. While these changes may collectively lead to the tropism switch, why does this occur in the majority of HIV-1 infections? How can random sequence evolution, given it’s blind nature, result in a predictable outcome? Could changes of flexibility in the gp120 V3 loop play a role in rewiring protein-protein interactions (from CCR5 to CXCR4) and therefore be a determinant of tropism?

The classical structure-function paradigm states that protein function requires a well-defined three-dimensional structure (23, 24). However, it has become clear that many functionally important proteins do not have well-defined structures (termed intrinsically disordered proteins) or have protein regions that lack structures (proteins with disordered regions)(23, 25-28). HIV-1’s gp120 has five highly variable (V1-V5) and five conserved (C1-C5) protein regions (29). The variable regions are normally missing in the solved X-ray crystal structures (V1/V2, V3, V4 and V5) of gp120 unless co-crystallized with other binding partners, presumably because they are intrinsically disordered. Disordered proteins or protein regions have been observed to be “sticky” and aggregate potentially forming promiscuous molecular interactions (25, 30, 31). Disordered regions are frequently found to undergo post-translational modifications, such as phosphorylation or N-linked glycosylation (24, 32). In addition, amino acid residues in proteins that give rise to structural disorder tend to be polar and charged (24, 25). Given the disordered nature of HIV-1’s V3 loop, the documented role of charge and glycosylation in the tropism switch, we hypothesize that intrinsic disorder of the V3 loop plays a critical role in determining coreceptor usage and switches in cell tropism: rewiring protein-protein interactions by changes in intrinsic disorder.

## Materials and methods

### Data and tropism prediction

We retrieved three data sets: (i) Full-length envelope protein (*Env*) sequences of HIV-1 subtype B with tropism information (R5, R5X4 or X4) from the Los Alamos National Laboratory HIV sequence database (http://www.hiv.lanl.gov/content/index). These sequences have annotated coreceptor usage based on biological data only, which is not from sequence-based bioinformatics prediction (http://www.hiv.lanl.gov/components/sequence/HIV/search/help.html#coreceptor). And, two intrapatient data sets: (ii) A dataset containing C2 to V5 regions of *Env*, as used by Shankarrapa(7). (iii) A dataset consisting of V1 to V3 regions of *Env*, as used by Mild (16, 33). We predicted tropism using Geno2pheno[454] with default parameters, and a false positive rate (FPR) cutoff of 5% was used to class sequences as CXCR4-using (‘true’) or R5 tropic (‘false’) (34). Note that geno2pheno does not distinguish between X4 and R5X4 viruses, this means that sequences classed as true could be either X4-or dual-tropic, which has important implications when interpreting the statistical test results.

### Comparison of protein disorder in the V3 loop of CCR5, CXCR4 and CCR5-CXCR4 tropic viruses

Bioinformatics methods for charactering probable disordered regions in amino acid sequences generally fall into two categories: (1) machine-learning methods trained on missing (presumed to be disordered) regions of experimentally solved X-ray crystallographic structures and these trained models used to predict disordered regions in protein sequences, or (2) physiochemical properties and pairwise amino acid interaction energies can be calculated to determine likely disordered regions of sequences. The former approach is prone to errors in the solved protein structures deposited in Protein Data Bank (http://www.rcsb.org/) (35). We therefore opted to use a method, IUPred (36) based on the latter approach to predict differences in intrinsic disorder regions between CCR5 and CXCR4 using virus in our datasets. A recent study comparing this method to other intrinsic disorder predictive methods concluded that IUPred makes the most accurate predictions (35, 37).

For each amino acid residue in a protein sequence, IUpred will report a disorder score from 0 to 1 ranging from complete order (0) to complete disorder (1). We use a cut-off 0.4 to indicate structural disorder (>=0.4) (37). Because IUPred uses a sliding window approach to calculate disorder over a range, we first performed disorder prediction on the full available envelope sequences in our three datasets, and then extracted the V3 region to analyze disorder within the V3 loop. We then group the extracted scores into two groups (CCR5-using and CXCR4-using) according to tropism prediction by geno2pheno, and compare these statistically using a nonparametric method in the R statistical package (38) (two sample comparisons: nonparametric Behrens-Fisher problem in paired data, http://cran.r-project.org/web/packages/nparcomp/index.html). We performed three statistical tests. First, the null hypothesis is *H*_0_ : *P*(*a*,*b*) 1 / 2 (b tends to be similar to a); the alternative hypothesis is *H*_*a*_ : *P*(*a*, *b*) < 1/ 2 or *P*(*a*, *b*) > 1/ 2(b tends to be smaller or larger than a). Second, the null hypothesis is *H*_0_ : *P*(*a*, *b*) □ 1/ 2 (b tends to be similar to/ larger than a); the alternative hypothesis is *H*_*a*_ : *P*(*a*, *b*) > 1/2 (b tends to be smaller than a). Finally, the null hypothesis is *H*_0_ : *P*(*a*, *b*) □ 1/ 2 (b tends to be similar to/smaller than a); the alternative hypothesis is *H*_*a*_ : *P*(*a*, *b*) > 1/ 2 (b tends to be larger than a).

## Results

First, we investigated whether there were statistically significant differences in structural disorder associated with coreceptor usage. We calculated the disorder tendency of amino acids in the full-length consensus envelope protein sequences (gp120 and gp41) of R5, R5X4 and X4 tropic viruses, respectively. This allowed us to obtain an overview of the extent of protein disorder in HIV-1 envelope proteins by tropism. The predicted disorder tendency within the three proteins groups coincides with regions missing from experimentally solved crystallized protein structures of gp120 (39) and gp41 (40)(fig. 1). For example, the electron densities of the longer intrinsically disordered regions such as V1/V2, V4 and V3 are difficult to obtain independently without co-crystallization with a stabilizing binding partner (protein regions longer than 30 amino acids, if solved, they are normally presented as flexible loops in the structure, see Fig 1A, B, C and D).

**Figure 1:**
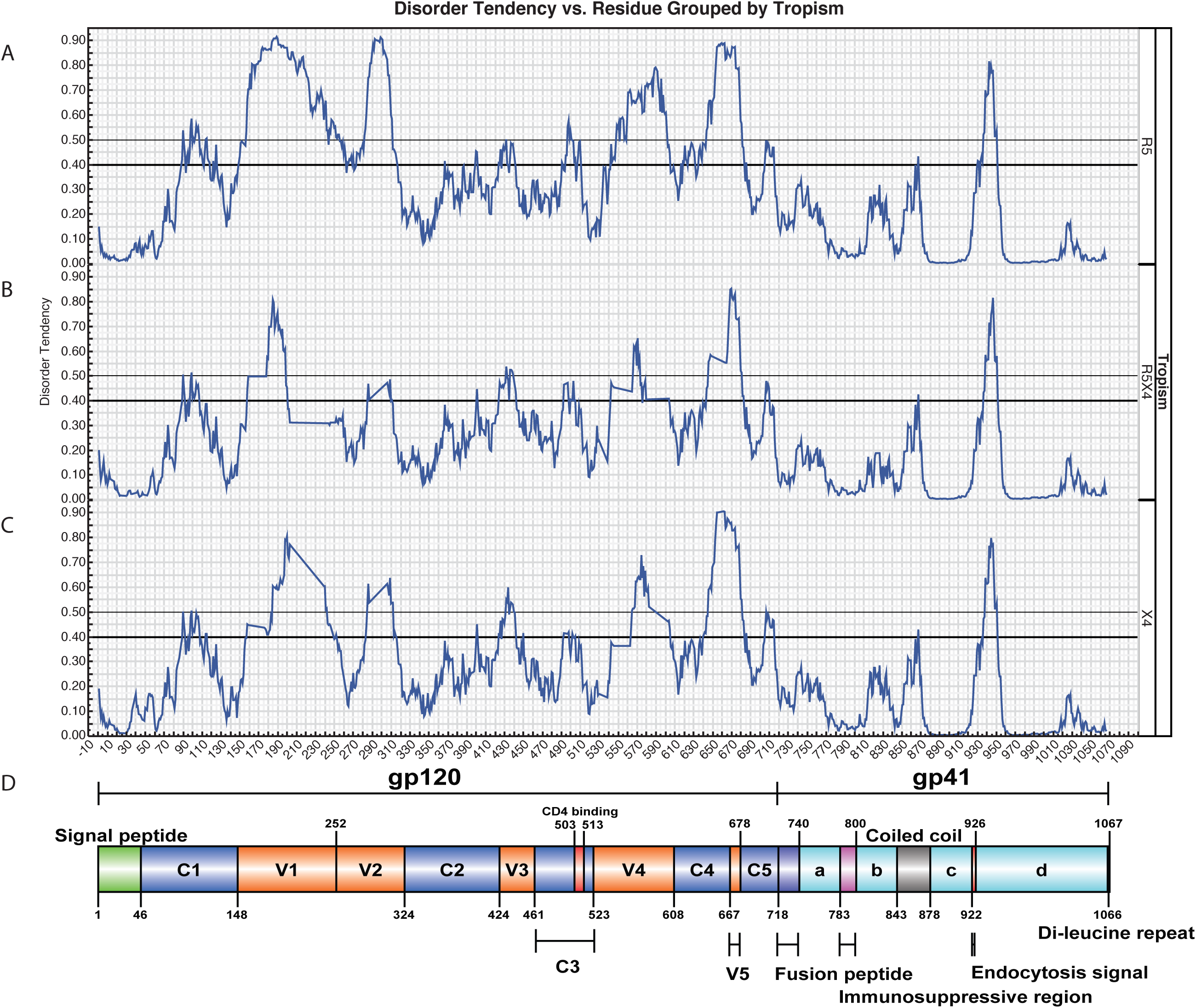
Plot of predicted structural disorder tendency of the consensus envelope protein. Disorder tendency for each amino acid is predicted by IUPred (0 represents complete order; 1 represents complete disorder; scores over 0.4 represent intrinsically disordered). Disorder tendency scores are plotted for consensus (A) R5, (B) R5X4 and (C) X4 envelope sequence, respectively. (D) The consensus protein sequences of R5, R5X4 and X4 are aligned to HXB2 envelope sequence (gp120 and gp41) for indicating the structure locations of the disordered residues. Gp120 and gp41 protein domains are numbered and color-coded for visualisation.

Next, we focus on the V3 loop to examine how structural disorder changes between coreceptor switches. We found that the disorder tendency of amino acids in the V3 loop of X4 virus was significantly higher than in R5 virus (*p*=2.35 × 10^−3^, fig 2) and R5X4 virus (*p*=1.87 × 10^−16^,fig 2), respectively. Interestingly, there was significantly higher structural disorder in R5 virus compared to R5X4 virus (*p*=7.49 × 10^−6^, fig 2). Thus, dual tropic R5X4 virus has the lowest V3 loop structural disorder tendency, compared to R5 and X4 virus. In sum, these results suggest that the V3 domain of X4 virus has significantly greater structural disorder tendency than that of R5 and R5X4 virus during coreceotor switch. Dual tropic R5X4 virus may therefore have a less flexible V3 domain than R5 and X4 virus, which may be more prone to neutralizing antibody binding. This is consistent with a previous study that showed dual tropic virus had lower fitness, and was therefore more sensitive to antibody inhibitors and neutralization (41).

**Fig 2.**
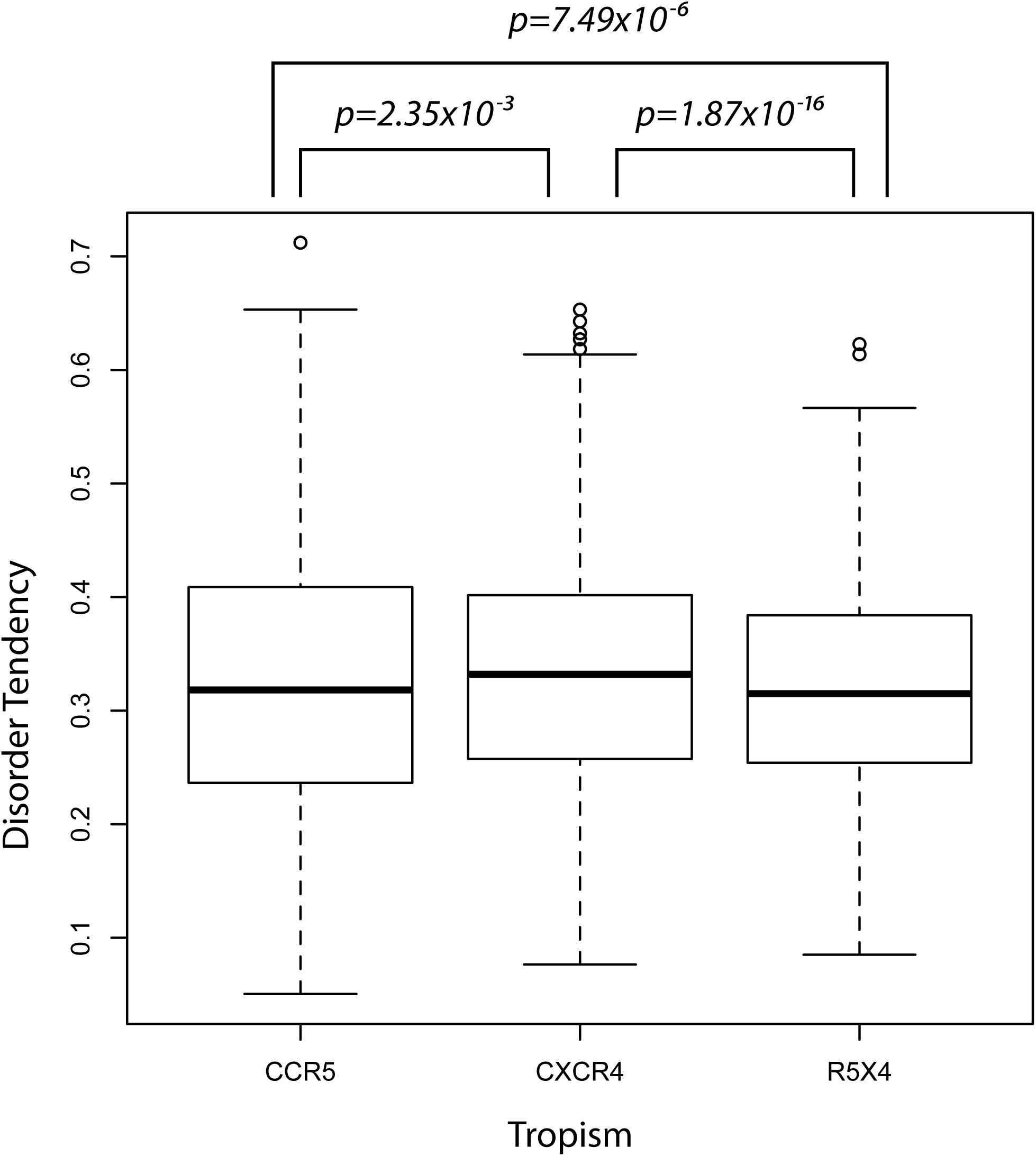
Boxplot and nonparametric comparisons of the V3 loop disorder tendency. Disorder scores are predicted by IUPred for all R5, R5X4 and X4 envelope sequences, respectively. Disorder tendency scores are compared by a nonparametric method described in Materials and Methods. The X4 V3 loop has significantly higher disorder scores than R5 (*p*= 2.35 × 10^−3^) and R5X4 (*p*= 1.87 × 10^−16^) virus, respectively. The V3 loop of R5 virus also has significantly higher disorder scores than R5X4 virus (*p*= 7.49 × 10^−6^).

To further test our hypothesis, we analyzed two data sets (Shankarappa et al. data and Mild et al. data) with longitudinal time points and multiple patients (7, 33). The Mild et al. dataset also had experimentally verified coreceptor calls. In the Shankarappa dataset, we first predicted coreceptor usage for all sequences at all visits across all nine patients (patient 1, 2, 3, 5, 6 7, 8, 9 and 11). We compared the structural disorder tendency between X4-capable and R5 virus at each visit that has coreceptor switch events in eight patients (patient 1, 2, 3, 5, 7, 8, 9 and 11). We showed the results for two patients (patient 2 and 9) that are representative of all eight patients, as suggested in Shankarappa et al. study (results for patient 1, 3, 5, 6, 7, 8 and 11 are reported in supplementary fig s1). In patient 2 we found that X4-capable V3 loop had significantly higher disorder tendency than R5 using V3 loop at visits 7 (p<0.001), 10 (p<0.05), 12 (p<0.001), 13 (p<0.001), 14 (p<0.05), 15 (p<0.001), 17 (p<0.001), 19 (p<0.001) and 23 (p<0.001) except at visit 16 (fig 3 patient 2). At visit 16 the R5 using V3 loop has significantly higher disorder tendency than the X4-capable V3 loop (the mean disorder tendency of the R5 using V3 loop is also higher than the X4-capable V3 loop), which may indicate the predicted X4-capable virus is actually dual tropic R5X4. For patient 9, it was only at visit 20 that the disorder tendency of the X4-cabable V3 loop is significantly higher than that of the R5 using V3 loop (p<0.001 fig 3 patient 9). At both visits 3 and 21 the R5 using V3 loop disorder tendency is significantly higher than the X4-cabable V3 loop indicating the predicted tropism may be dual tropic R5X4. However, at both visits 17 and 22 there is no significant difference of disorder tendency during corecetor switch (p>0.05 fig 3 patient 9), which may suggest the prediction of tropism is false positive and they have the same tropism.

**Fig 3.**
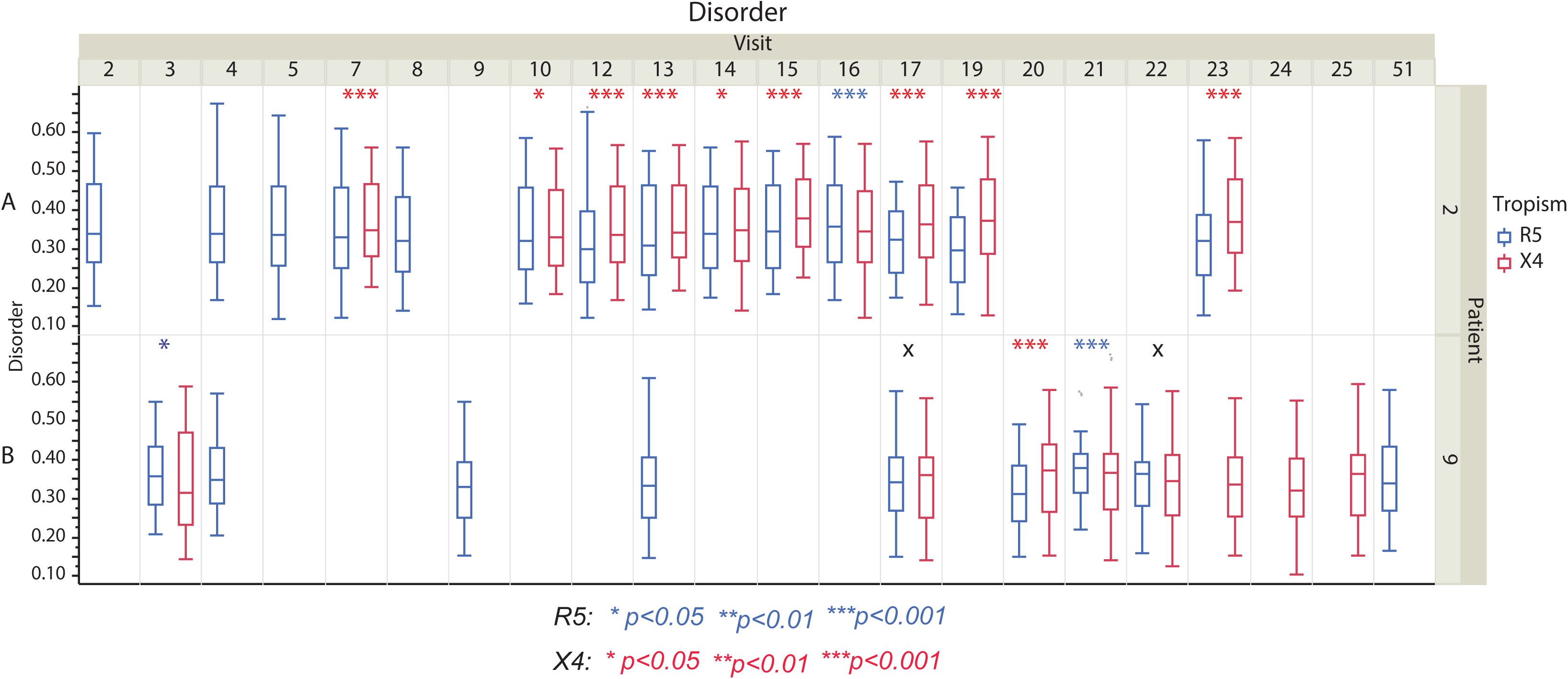
Boxplot and nonparametric comparisons of the V3 loop disorder tendency between R5 and X4 viruses from Shankarappa et al. data (7). Boxplot and statistical comparisons are shown for patient 2 (A) and 9 (B), respectively (\**p*<0.05, \*\**p*<0.01, *\*\*\*p*<0.001, asterisks colored with blue and red represent results for R5 and X4 virus, respectively; X means *p*>0.05).

In the Mild et al. data we first predicted the coreceptor usage for all sequences at each time point within patients. Patients who are predicted to have coreceptor switch events are consistent with the experiment verification except 25 months, 45 months, 10 months in patient 2239, 2242 and 2282, respectively (patient 2239, 2242 and 2282 are included in our study while patient 1865 was omitted due to dual tropic R3X4 virus). We then calculated the disorder tendency in R5 and X4 virus (fig 4), respectively. We finally compared the disorder tendency between R5 and X4 virus at each time point of all three patients that have coreceptor switch events. In patient 2239 we find that there are three time points that have coreceptor switch events (time points 25 months, 68 months and 88 months). At two of the three time points the X4-capable V3 loop has significantly higher disorder tendency than the R5 using V3 loop (p<0.001). In patient 2242 we find three time points that have coreceptor switch events (time points 45 months, 84 months and 85 months). However, none of the three switch events involves any significant change of V3 loop disorder tendency (p>0.05, fig 4 patient 2242). In patient 2282 we find 5 time points that have coreceptor switch events (time points 10 months, 47 months, 62 months, 63 months and 70 months, fig 4 patient 2282). Interestingly, all X4 V3 loops have significantly higher structural disorder tendency than R5 using V3 loops (p<0.001, fig 4 patient 2282). These results are compatible with our hypothesis: Firstly the X4-capable virus had significantly higher structural disorder than the R5 tropic virus (time points 25 months and 88 months in patient 2239; time points 10 months, 47 months, 62 months 63 months and 70 months in patient 2282, fig 4). Secondly, the structural disorder is not significantly different between R5 and X4 tropic virus (time points 45 months, 84 months and 85 months in patient 2242, fig 4). This is not consistent with our hypothesis, which may suggest that the predicted tropism is due to false prediction and therefore we may compare two groups of virus with the same tropism. Moreover, the mean disorder tendency in each comparison was also consistent with our hypothesis, although some comparisons did not reach statistical significance (except time point 45 months in patient 2242).

**Fig 4.**
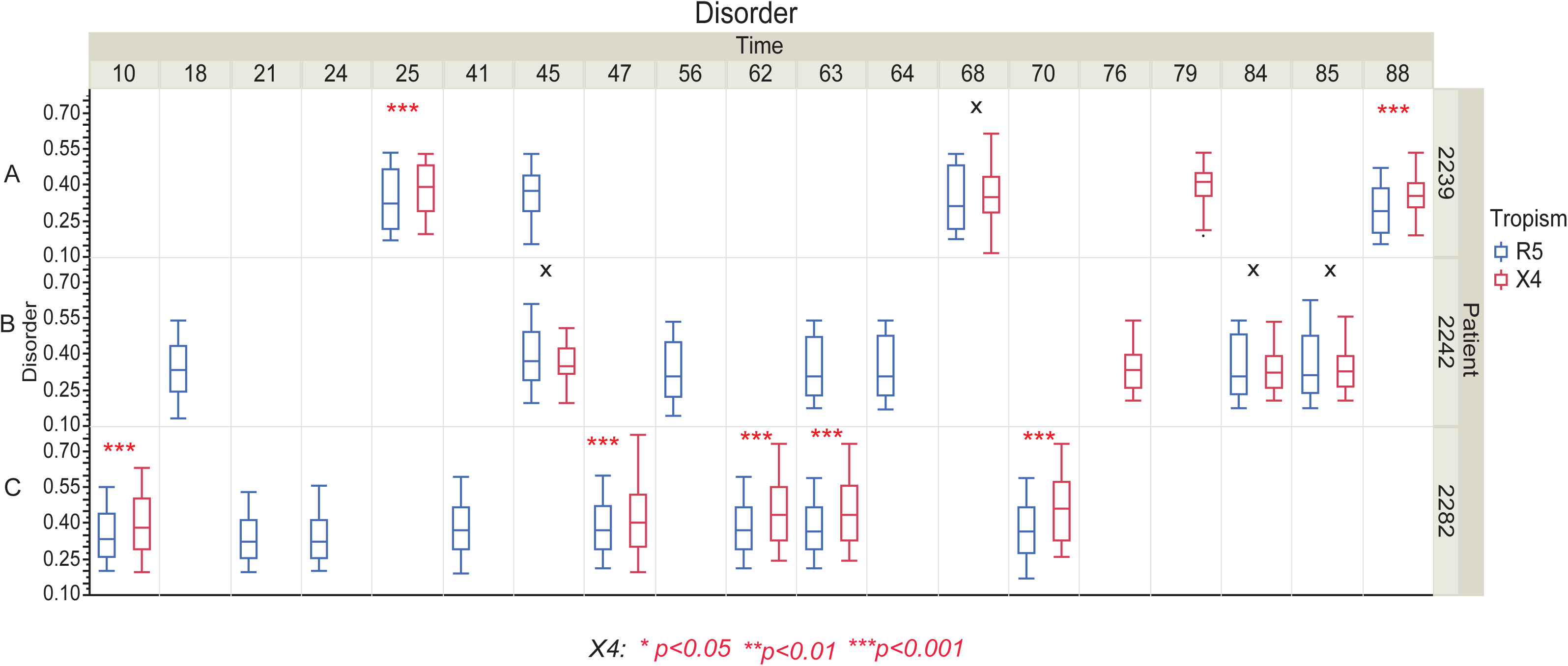
Boxplot and nonparametric comparisons of the V3 loop disorder tendency between R5 and X4 viruses from Mild et al. data (33). Boxplot and statistical comparisons are shown for patient 2239 (A), 2242 (B) and 2282 (C), respectively (\**p*<0.05, \*\**p*<0.01, *\*\*\*p*<0.001, asterisks colored with red represent results for X4 virus; X means *p*>0.05).

## Discussion

In this study we have used a nonparametric statistical approach to investigate if there are distinct patterns of predicted structural disorder in the V3 domain of HIV-1 envelope protein between R5, R5X4 and X4 tropic viruses. Strikingly, there seems to be an increase of structural disorder tendency of V3 domain from R5/R5X4 tropic virus to X4 tropic virus, which indicates that structural disorder (flexibility) of the V3 loop plays a role in HIV-1 cell tropism.

Our statistical comparisons of the predicted disorder tendency between disordered V3 loops capable of using different chemokine receptors (CCR5 and/or CXCR4) are consistent with our hypothesis that 1) the V3 loop of X4 using virus has significantly higher disorder tendency than that of R5 virus; and 2) the V3 loop of X4 and R5 virus has significantly higher disorder than the dual tropic (R5X4) virus. As protein disorder is a source of novel protein-protein interactions (23, 24, 30, 35, 42), such a significant shift of protein disorder tendency is presumably contributing to rewiring of protein-protein interactions (42).

In terms of a mechanism for the switch, intrinsically disordered regions are relatively unconstrained resulting in the accumulation of neutral or nearly neutral residue changes in line with the neutral theory of molecular evolution (43, 44). Random changes may lead to some affinity for the CXCR4 coreceptor, i.e., dual tropic intermediate viruses (41) which selection may then act upon resulting in X4 virus.

The higher disorder tendency of the X4 virus V3 loop suggests it may be ‘stickier’ and able to use CXCR4 coreceptor more efficiently and therefore cause further infection of immune cells that express CXCR4. Alternatively, increased disorder tendency of the V3 loop may make the virus more promiscuous in binding to immune cell coreceptors for cell entry (e.g., not just CXCR4) (30, 45).

We speculate that that structural disorder of the V3 loop contributes to function by being stabilized by secondary interactions leading to coreceptor switch, which could be another protein (e.g., CXCR4 or other chemokine receptors (45)) or posttranslational modifications such as N-linked glycosylations as suggested in other studies (13-20, 22). Evidence from several previous studies supports our hypothesis. First, non-switching viral populations (remain R5 using) have increased N-linked glycosylation sites, which has a stabilization role (17, 19-22). Second, molecular-dynamics simulation study also suggests that X4 binding stabilizes the V3 loop (46). Third, many previous studies demonstrate that increased charge in the V3 loop play a significant role in coreceptor switch (13, 16, 17, 47, 48). It is well known that charged residues promote structural disorder (24), which leads to increased disorder in X4 using V3 loop. However, if the V3 loop with increased disorder can be stabilized by increased N-glycans or a combination of N-glycans and other secondary interactions the virus could remain R5 tropic (13-18, 22).

In conclusion, understanding the evolutionary mechanisms that lead to the fixation of virus protein structural disorder may hold the key to understand viral host adaptation and develop novel intervention strategies. Here the HIV-1 structural disorder mediated coreceptor switch provides a unique model to study change in specificity of a protein-protein interaction, which provides a mechanistic understanding of HIV-1 cell tropism. Future research aiming at understanding viral host adaptation could benefit from elucidating the molecular evolutionary mechanisms that lead to the fixation of structural disorder promoting mutations in the virus-host protein interaction networks, as these mutations may be key to host adaptations and cross-species transmission.

## Acknowledgements

We would like to thank Simon Lovell for helpful discussion, and Frank Konietschke and Marius Placzek for providing an updated version of the R Nparcomp package. FF was supported by a BBSRC studentship to DLR. XJ was supported by the MRC (G1001806/1) and WT (097820/Z/11/A) to DLR and by the Issac Newton Trust and WT (PCJW/GAAB) funding to John Welch, University of Cambridge.

## Author contributions

XJ and DLR conceived and designed the study. XJ performed the research. XJ, DLR and FF analysed the data. XJ, DLR and FF wrote the paper. All authors commented on and approved a final version of paper.

